# Genetic interplay between type II topoisomerase enzymes and chromosomal *ccdAB* toxin-antitoxin in *E. coli*

**DOI:** 10.1101/2021.09.24.461737

**Authors:** Jay W. Kim, Vincent Blay, Portia Mira, Miriam Barlow, Manel Camps

## Abstract

Fluoroquinolones are one of the most widely used class of antibiotics. They target two type II topoisomerase enzymes: gyrase and topoisomerase IV. Resistance to these drugs, which is largely caused by mutations in their target enzymes, is on the rise and becoming a serious public health risk. In this work, we analyze the sequences of 352 extraintestinal *E. coli* clinical isolates to gain insights into the selective pressures shaping the type II topoisomerase mutation landscape in *E. coli*. We identify both Quinolone Resistance-Determining Region (QRDR) and non-QRDR mutations, outline their mutation trajectories, and show that they are likely driven by different selective pressures. We confirm that ciprofloxacin resistance is specifically and strongly associated with QRDR mutations. By contrast, non-QRDR mutations are associated with the presence of the chromosomal version of *ccdAB*, a toxin-antitoxin operon, where the toxin CcdB is known to target gyrase. We also find that *ccdAB* and the evolution of QRDR mutation trajectories are partially incompatible. Finally, we identify partial deletions in CcdB and additional mutations that likely facilitate the compatibility between the presence of the *ccdAB* operon and QRDR mutations. These “permissive” mutations are all found in ParC (a topoisomerase IV subunit). This, and the fact that CcdB-selected mutations frequently map to topoisomerase IV, strongly suggests that this enzyme (in addition to gyrase) is likely a target for the toxin CcdB in *E. coli*, although an indirect effect on global supercoiling cannot be excluded. This work opens the door for the use of the presence of *ccdB* and of the proposed permissive mutations in the genome as genetic markers to assess the risk of quinolone resistance evolution and implies that certain strains may be genetically more refractory to evolving quinolone resistance through mutations in target enzymes.

## Introduction

Gyrase and topoisomerase IV are essential enzymes in most pathogenic bacteria. These enzymes belong to the family of type II topoisomerases: they regulate topology and torsional stress of DNA by catalyzing a double-strand break and passing a double strand through the break (1). Both enzymes form heterotetramer (A_2_B_2_) enzyme complexes, with “A” representing the DNA strand-passing subunit, and “B” representing the ATPase subunits (2,3). The two enzyme complexes exhibit a high degree of structural homology and are encoded in *E. coli* by *gyrA* (A subunit) and *gyrB* (B subunit) genes in the case of the gyrase, and by *parC* (A subunit) and *parE* (B subunit) in the case of topoisomerase IV (2,3).

Quinolones are drugs that target gyrase and topoisomerase IV (4) and represent one of the most widely used classes of antibiotics used to treat bacterial infections (5). Specifically, quinolones stabilize a catalytic intermediate that forms once the DNA strand is broken by the nucleophilic attack of a tyrosine in the enzyme active site (6). These stable “cleaved complexes” lead to bacterial cell death through complex mechanisms (7). Whether the main target of quinolones is DNA gyrase or topoisomerase IV seems species- and compound-specific (8), although highly resistant organisms tend to contain resistance-conferring mutations in both enzymes (5,8-10).

The rates of quinolone resistance are increasing rapidly in recent years (11-13) and therefore an understanding of this evolutionary process becomes crucial to ensure global health. Quinolone resistance can arise by several mechanisms. One of the most prominent mechanisms is a reduction in the binding affinity of the type II topoisomerase-DNA complex for quinolones through mutations in the binding site (14). These mutations are typically located in an area known as the Quinolone Resistance-Determining Region (QRDR) (15,16). The most frequent QRDR mutations in *E. coli* occur at positions S83 and D87 of GyrA and S80 or E84 of ParC. ParE positions L416 and S458 also represent QRDR mutations contributing to fluoroquinolone resistance (10). A second mechanism is mediated by quinolone resistance (*qnr*) genes. These genes encode proteins that bind to gyrase and topoisomerase IV and prevent quinolone binding (17-20). Other mechanisms involve lowering the drug concentration by upregulating active efflux transporters, downregulating porin diffusion channels that allow its passive influx (15,21,22), or chemically modifying the drug by an aminoglycoside acetyltransferase (AAC(6’)-lb-cr) (23). Most of the mechanisms discussed above are chromosomal, but *qnr* genes and *aac(6’)-Ib-cr* are typically plasmid-borne.

Here, we study the sequences and fluoroquinolone resistance phenotypes of 352 clinical *E. coli* isolates from two US West Coast hospitals to learn about the evolution of fluoroquinolone resistance. In this dataset, the “ciprofloxacin-resistant” class is tightly associated with Quinolone Resistance-Determining Region (QRDR) mutations in type II topoisomerase genes. We also find many non-QRDR mutations both in gyrase and in topoisomerase IV whose evolution appears to be driven by a different, older selection, which we trace to a chromosomal version of the toxin CcdB. This toxin appears to be largely incompatible with QRDR mutations unless partially deleted or in the presence of certain permissive mutations. Thus, our results provide strong genetic evidence for an interplay between chromosomal CcdB toxin and type II topoisomerase genes. These observations shed new light into the evolution of fluoroquinolone resistance and open the door for the use of *ccdB* and of permissive mutations as genetic markers of risk of quinolone resistance evolution.

## Results and Discussion

The present study is based on the analysis of two independent whole-genome datasets that are accompanied by results from antibiotic susceptibility testing against a panel of antibiotics, including the fluoroquinolone ciprofloxacin. The joint dataset, USWest352, combines 233 genomic sequences previously reported from University of Washington hospital (24) and 119 new genomic sequences from extended-spectrum β-lactamase resistant isolates of Dignity Health Merced Medical Center generated for this study. In this joint dataset, 48% of the isolates displayed ciprofloxacin resistance, providing the balanced representation of ciprofloxacin-sensitive and -resistant samples necessary for this study. Specific details about the two datasets can be found in **Methods**.

USWest352 contains a total of 85 different sequence types (STs). STs result from clustering multiple strains based on the similarity of genetic markers throughout the genome (see **Methods**) (25). The full distribution of sequence types in our dataset is listed in **Dataset S1** and a summary of the strain composition is provided in **Table S1**. We see an overrepresentation of epidemic STs (i.e., sequence types highly prevalent across the globe), with four STs accounting for 49% of the samples. To correct for the effects of the observed clonal expansion of these and other strains, samples were grouped in sequence types (STs) prior to analysis.

The genomes were analyzed for known fluoroquinolone resistance determinants. These include the six QRDR positions known to be associated with fluoroquinolone resistance (10,15); plasmid-encoded, DNA-binding protective factors (qnr); and the aminoglycoside acetyl transferase gene *aac(6’)-lb-cr* (see **Methods**). The results are listed in **Dataset S1**. The QRDR mutation tallies are listed in the top panel of **Table 1** along with the variation in the amino acid substitutions represented at these positions. Our observations generally agree with previous studies, and ParC-E84Q appears to be an amino acid substitution that has not been reported before.

**Table 1.**
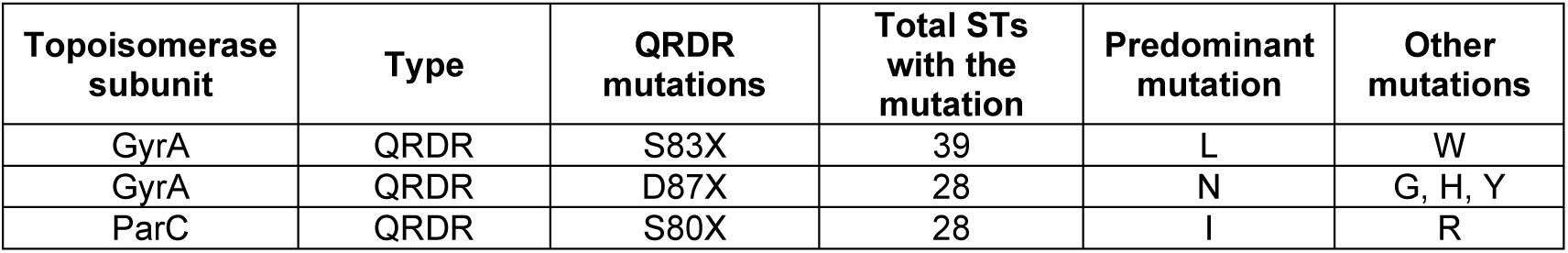
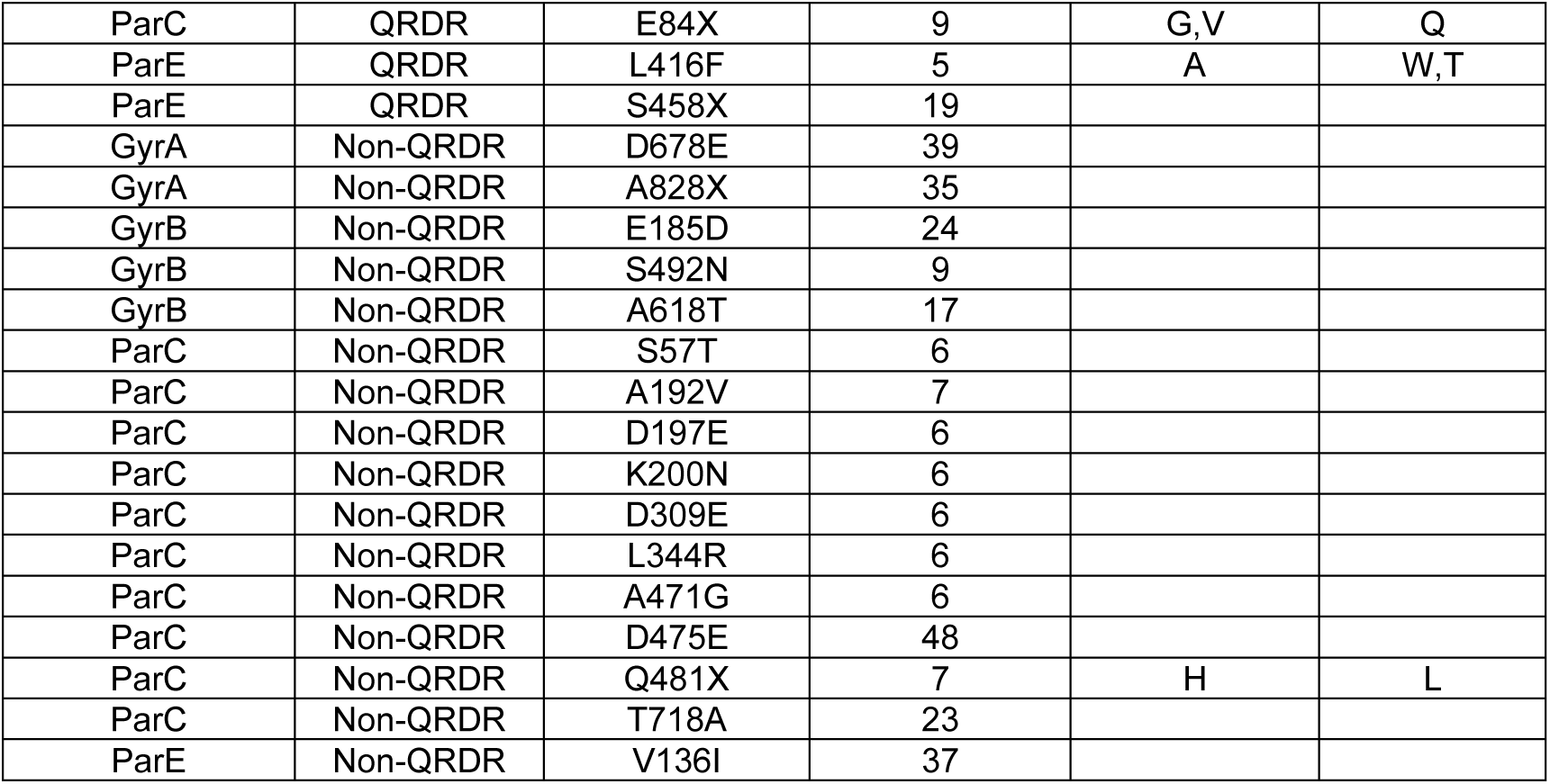
Gyrase and topoisomerase IV mutations observed in five or more STs per position in dataset USWest352. Known QRDR mutations and non-QRDR mutations are reported, and additional information is given for polymorphic sites.

| Topoisomerase subunit | Type | QRDR mutations | Total STs with the mutation | Predominant mutation | Other mutations |
| --- | --- | --- | --- | --- | --- |
| GyrA | QRDR | S83X | 39 | L | W |
| GyrA | QRDR | D87X | 28 | N | G, H, Y |
| ParC | QRDR | S80X | 28 | I | R |
| ParC | QRDR | E84X | 9 | G,V | Q |
| ParE | QRDR | L416F | 5 | A | W,T |
| ParE | QRDR | S458X | 19 |  |  |
| GyrA | Non-QRDR | D678E | 39 |  |  |
| GyrA | Non-QRDR | A828X | 35 |  |  |
| GyrB | Non-QRDR | E185D | 24 |  |  |
| GyrB | Non-QRDR | S492N | 9 |  |  |
| GyrB | Non-QRDR | A618T | 17 |  |  |
| ParC | Non-QRDR | S57T | 6 |  |  |
| ParC | Non-QRDR | A192V | 7 |  |  |
| ParC | Non-QRDR | D197E | 6 |  |  |
| ParC | Non-QRDR | K200N | 6 |  |  |
| ParC | Non-QRDR | D309E | 6 |  |  |
| ParC | Non-QRDR | L344R | 6 |  |  |
| ParC | Non-QRDR | A471G | 6 |  |  |
| ParC | Non-QRDR | D475E | 48 |  |  |
| ParC | Non-QRDR | Q481X | 7 | H | L |
| ParC | Non-QRDR | T718A | 23 |  |  |
| ParE | Non-QRDR | V136I | 37 |  |  |

In our dataset, complex QRDR mutants always contain mutations at GyrA S83, making this mutation a gateway to different quinolone-driven adaptive trajectories. This is consistent with previous results indicating that gyrase is the preferred target of fluoroquinolones in Gram negative bacteria (5). 72% of S83X mutant samples also had mutations at positions S80 of ParC. Secondary targets for fluoroquinolones (topoisomerase IV in this case) acquire mutations once the first-step mutation is fixed and the affinity for the secondary target limits the increase in resistance (26).

Mutations ParC-S80X and GyrA-D87N are almost always observed together. Downstream of the GyrA S83X, D87N and ParC S80X triplet, we see three largely mutually exclusive trajectories (**Fig. 1**). The first one has an additional mutation in L416 of ParE (to F). The second one has additional amino acid substitutions in position E84 of ParC (to G,V, or Q). The third one has mutations at position S458 in *parE* (to A,W,T). The observed trend toward mutual exclusivity suggests that these three mutations could be playing redundant roles in enhancing fluoroquinolone resistance.

**Fig. 1.**
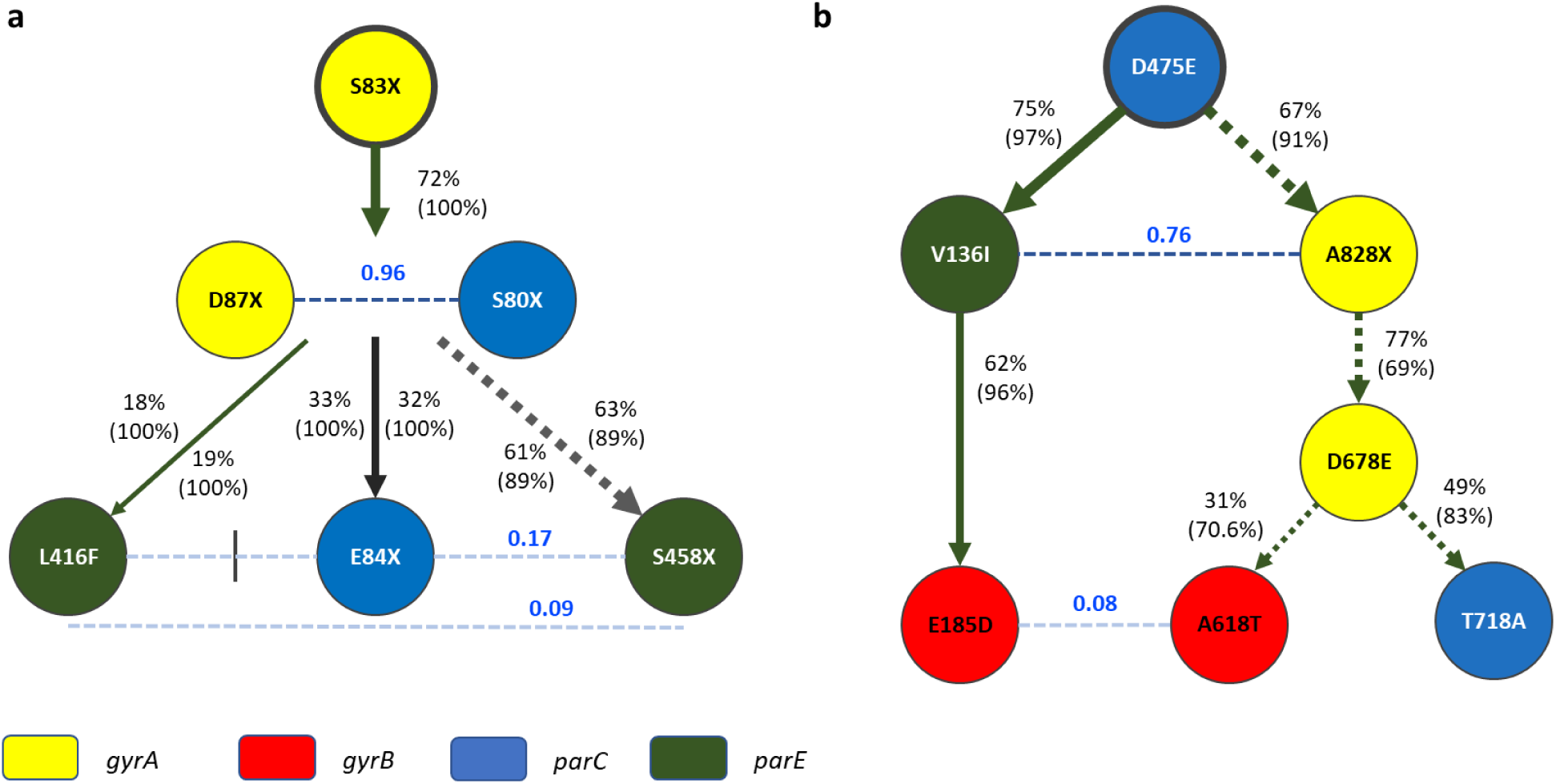
Main type II topoisomerase mutation pathways observed in USWest352. **a** QRDR pathway. **b** Non-QRDR pathways. Positions that are frequently mutated in these pathways are shown as nodes. Nodes are color-coded according to gyrase or topoisomerase IV subunit, with GyrA in yellow, GyrB in red, ParC in blue, and ParE in dark green. Percentages represent the fraction of the downstream mutation contained in mutants containing the upstream mutation. Percentages in parenthesis represent the fraction of samples with the downstream mutation that also have the upstream mutation. When this percentage is above 95% the arrow is solid. When less than 95%, the arrow is dotted. Blue numbers on dotted lines represent Jaccard indexes between the two nodes joined by the line. The Jaccard index (JI) measures the co-occurrence of two mutations and ranges from 0 (mutual exclusion) to 1 (perfect co-occurrence) (see **Methods**).

### Non-QRDR mutations define an alternative evolutionary pathway for type II topoisomerases

Besides QRDR mutations, samples in USWest352 also had variants in many additional positions in type II topoisomerase genes. We refer to these as “non-QRDR” mutations. All non-QRDR positions present in five or more STs are tallied in **Table 1**. We investigated whether non-QRDR mutations represent one or multiple adaptive trajectories and how these trajectories relate to QRDR adaptation. To this end, the Jaccard indexes between all mutation pairs were computed. The Jaccard index (JI) measures the co-occurrence of two mutations (or other binary features), and ranges from 0 (mutual exclusion) to 1 (perfect co-occurrence) (see Methods). **Fig. 2** shows the JI in the form of a heatmap for all pairwise combinations of mutations present in five or more STs.

**Fig. 2.**
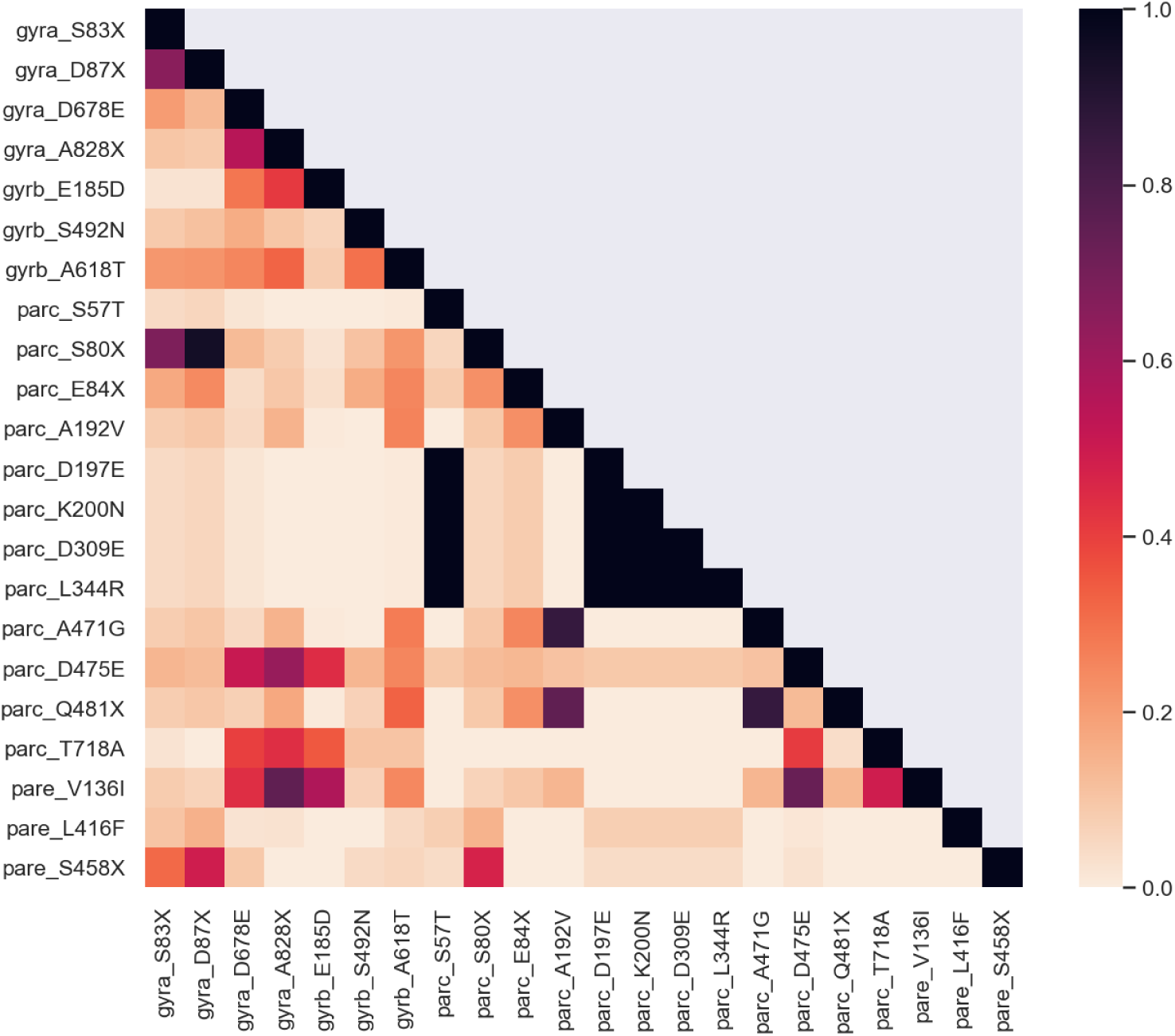
Jaccard indexes between commonly occurring mutations in type II topoisomerases. Jaccard indexes for all gyrase and topoisomerase IV mutations at positions observed to have mutated in at least five STs are presented as a heatmap, with values closer to 1 (dark) indicating a trend toward co-occurrence and values closer to zero (light) indicating a trend toward mutual exclusion.

Non-QRDR mutation trajectories are summarized in **Fig 1b**, although they cannot be as clearly delineated as in the case of QRDR mutations. ParC-D475E is present in most pairwise combinations containing the other non-QRDR mutations. The GyrA-A828X, ParC-D475E, and ParE-V136I triplet is the most frequent combination of non-QRDR mutations. Three downstream trajectories can be derived from this triplet. Among these downstream trajectories, we also noted that GyrB A618T and E185D mutations show strong mutual exclusion, suggesting either functional redundancy or functional incompatibility.

In ParC, the Jaccard index also shows blocks of mutations that co-occur frequently. One of these blocks includes five mutations: [S57T D197E K200N D309E L344R]. The other block involves three mutations: [A192V A471G Q481H/L]. In both cases, the exact combination of ParC mutations appears in six different STs, although A192V and Q481H/L are also seen once outside this combination.

### Non-QRDR and QRDR mutations are driven by different selective pressures

To look at the distribution of QRDR and non-QRDR mutations in the population, we generated a phylogeny based on nucleotide-level variation across whole genomes in USWest352 and visualized the phylogenetic relationship as a neighbor-joining (NJ) tree (see **Methods** and **Fig. 3**).

**Fig. 3.**
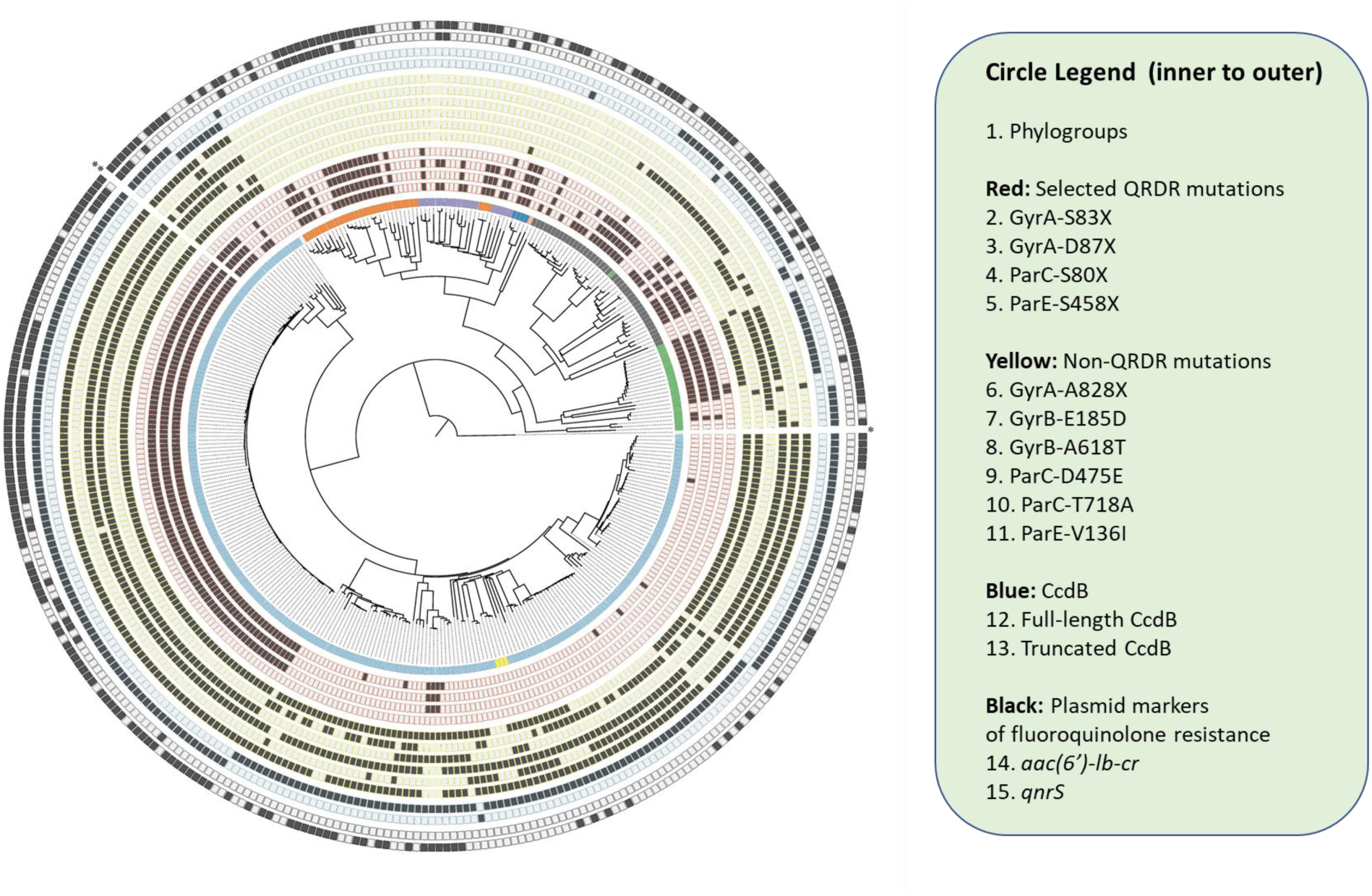
Visualization of the phylogenetic distance between 352 extraintestinal *E. coli* isolates obtained from two hospitals in the US West Coast. The phylogenetic relationship between the samples is represented as a neighbor-joining tree, computed based on pairwise comparisons of 537,420 genomic SNPs. The tree was constructed from an all-by-all pairwise distance matrix containing the number of nucleotide positions differing between each pair of genomes (27). The squares represent the presence of selected mutations or genes. Panel 1: Phylotype A (orange), B1 (purple), B2 (light blue), C (dark blue), D (dark grey), E (light red), F (green), U (yellow). Panels 2-5 (outlined in blue) selected QRDR mutations GyrA-S83X and -D87X, ParC S80X, ParE S458X. Panels 6-13 (outlined in yellow) Selected non-QRDR mutations: GyrA A828X, GyrB E185D and A618T, ParC 475E and T718A, and ParE V136I. Panel 14-15: CcdB (outlined in X): full-sized Ccdb, and truncated CcdB. Panels 16-17: Plasmid markers of fluoroquinolone resistance (outlined in Y) *aac(6’)-lb-cr, qnrS*.

QRDR mutations are found in a wide range of phylotypes. The clustered distribution of these QRDR mutations is consistent with clonal expansions, and the relatively small size of the clusters suggests that these clonal expansions are recent. To confirm this, we calculated an intra-ST spread index for QRDR mutations (see **Methods**). The results shown in **Fig. 4**. are consistent with a recent selection, with the gateway mutation GyrA S83X showing a intra-ST spread index of only 0.6, and the rest of the QRDR mutations ranging between 0.42 and 0.78 (**Fig. 4**, red bars). This observation is consistent with the frequent use of quinolones in the clinic (4,12), which is probably more recent than the last common ancestor for most STs, and with reports indicating that QRDR mutations have limited or no fitness costs and may even positive influence fitness (5,28), possibly through global changes in supercoiling (31,32).

**Fig. 4.**
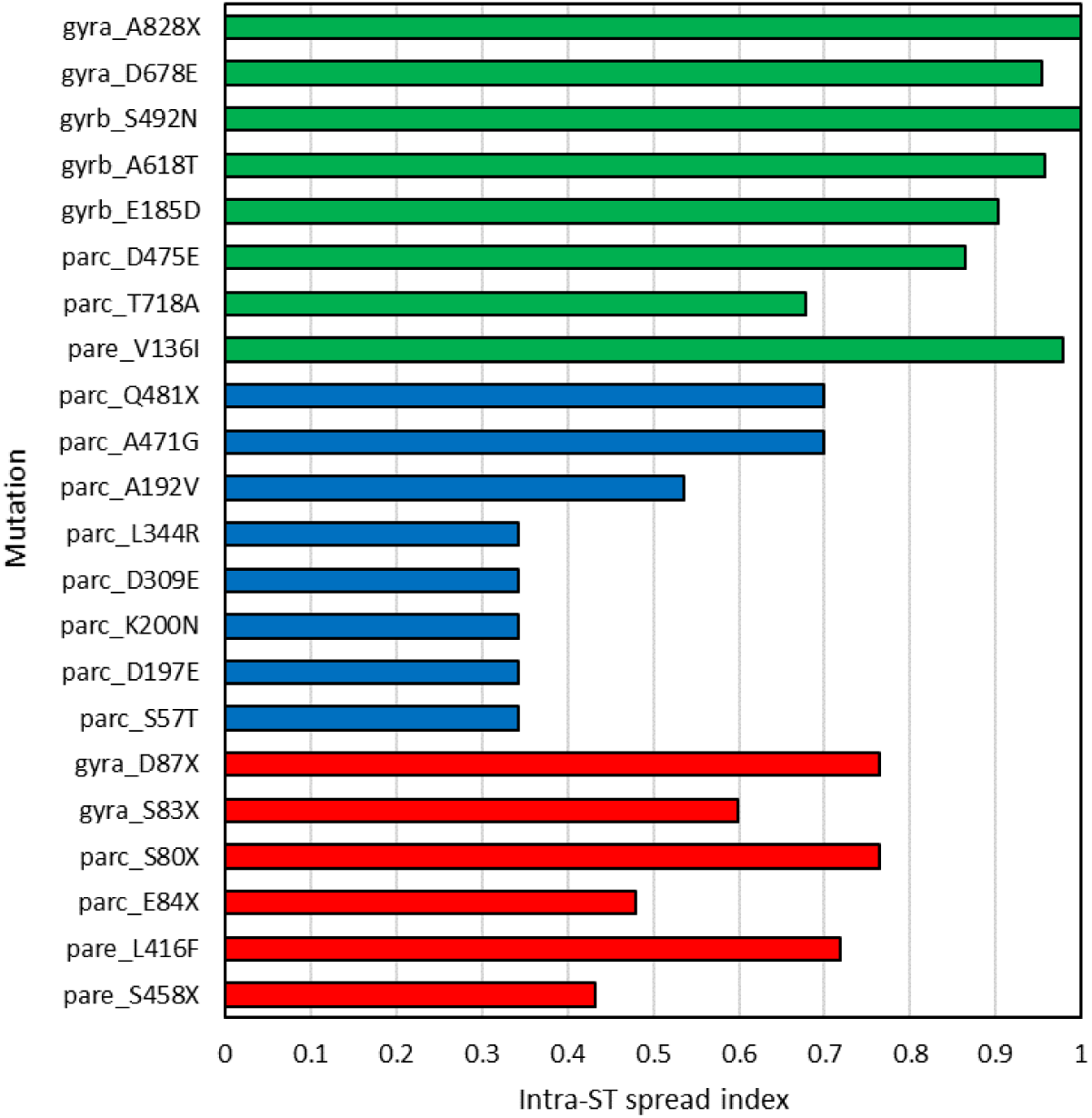
Intra-ST spread index for selected mutations in type II topoisomerase enzymes. The Intra-ST spread index measures the consistent presence of a mutation within sequence types, for sequence types in which at least three individuals contain the mutation. See **Methods** for details. The mutations are colored based on non-QRDR (green), permissive (blue), and QRDR (red).

The inconsistent presence of QRDR mutations within STs (as indicated by the intra-ST spread index in **Fig4**) can also be caused by reversions. This scenario cannot be ruled out given the negative impact some QRDR mutations on gyrase activity in vitro (29,30), although reversions are stochastically much more improbable than other compensatory mechanisms (28,31).

Non-QRDR mutations, by contrast, are completely absent in the sector of the cladogram that includes phylotypes A, B1, C, and E (**Fig. 3**). The putative clonal expansions that we observe for non-QRDR mutations are much larger in size, but rarely leading to a complete fixation of the mutation (**Fig. 3**, panels 6-11). Relative to QRDR mutations, non-QRDR mutations are more consistently present within STs, as indicated by higher intra-ST spread index (**Fig. 4**, compare green with red bars). Taken together, these observations suggest that non-QRDR mutations are the result of early fixation combined with additional positive and negative selections leading to inconsistent mutation trajectories.

The cladogram also confirms that the two colinear combinations of ParC mutations detected by the Jaccard indexes (**Fig. 2**) originated multiple times independently, since these combinations are seen in multiple, clearly separated sections of the cladogram. Specifically, the [S57T D197E K200N D309E L344R] and [A192V A471G Q481X] combinations are both seen in five separate sections of the cladogram (**Fig. S1**). In the case of [S57T D197E K200N D309E L344R], we even find this combination in two different phylotypes (D and B2). This combination also has a low intra-ST spread index (0.33), suggesting that these mutations are under recent positive or negative co-selection. The presentation of these mutations is completely binary, that is, either see all five mutations or none are observed, which points to a functional/synergistic interaction linking these mutations together. The other combination [A192V A471G Q481X] has a higher intra-ST spread index, but also shows a largely binary distribution. **Fig. S2** shows the location of these groups of co-linear mutations on the secondary structure of a topoisomerase dimer. Interestingly, these two groups of mutations fall on the opposite side of the enzyme active site.

### Association between non-QRDR mutations and chromosomal ccdAB

To try to identify a selection driving the fixation of non-QRDR mutations, we looked at another molecule known to target gyrase, namely the toxin CcdB. This toxin inhibits gyrase by a mechanism akin to that of fluoroquinolones, involving a direct interaction with gyrase and the stabilization of DNA-gyrase complexes (32). The toxin CcdB belongs to a toxin-antitoxin system (TAS). TAS consist of small genetic modules encoding a toxin that induces cell death or growth arrest and an antitoxin counteracting the activity of the toxin (33,34). CcdB’s corresponding antitoxin is CcdA. This antitoxin removes CcdB from poisoned complexes through interactions with CcdA’s folded C-terminus (35).

We hypothesized that, despite the presence of the antitoxin, CcdB toxin may place a selective pressure on the gyrase that is distinct from fluoroquinolone selection. To test the hypothesis that non-QRDR mutations are associated with the presence of *ccdAB*, we identified plasmid and chromosomal forms of *ccdAB* in our samples (listed in **Dataset S1**). The location of this TAS in the chromosome was confirmed using two independent methods (see **Methods** and Fig. **S3**).

We noted that distribution of chromosomal *ccdAB* in the cladogram closely tracks the presence of non-QRDR mutations (**Fig. 3**). Like non-QRDR mutations, *ccdAB* is absent from samples belonging to phylotypes A, B1, C, and E, while it is present in most other samples. Moreover, *ccdAB* co-occurs nearly always with ParC-D475E, the gateway mutation for non-QRDR mutation trajectories. The distribution of plasmid *ccdAB*, by contrast, is more similar to that of other chromosomal markers and does not correlate with the presence of non-QRDR mutations (not shown).

### Genetic interactions between mutations and GyrA S83X or chromosomal ccdAB

We next looked at possible genetic interactions between individual type II topoisomerase mutations and GyrA-S83X mutation (as an indicator of quinolone selection) and *ccdAB* (as a possible selection for non-QRDR mutations) (**Fig. 5**)

**Fig. 5.**
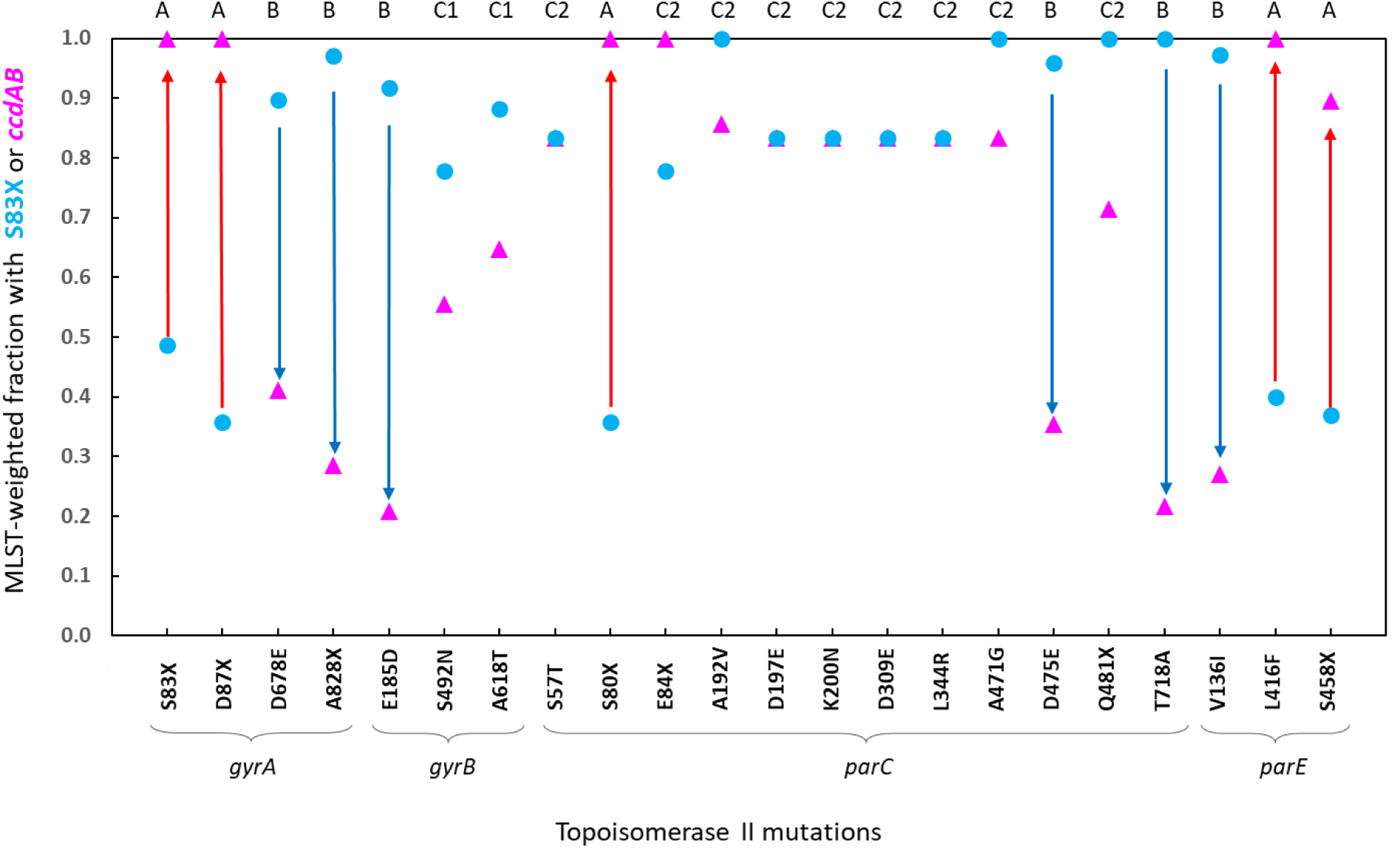
ST-weighted fractions of mutants containing S83X and of mutants containing CcdB. The presence of the QRDR gateway mutation S83X was quantified as a fraction of the total number of sequenced STs containing the mutation listed on the X-axis (blue dots). The presence of the *ccdAB* module was also quantified in the same way as a fraction of total STs (red dots). Blue arrows indicate a strong bias for S83X mutations relative to *ccdAB*. Green arrows indicate a strong bias for *ccdAB*. The mutations included are listed on the X-axis, and the subunit they belong to is indicated below. The proposed classification into four groups (A, B, C1 and C2) for each mutation is indicated at the top.

Based on these interactions (epistasis) we classified type II topoisomerase mutations into the following categories: mutations that frequently co-occur with GyrA-S83X and rarely with *ccdAB* (group A); mutations that rarely co-occur with GyrA-S83X, while frequently co-occurring with c*cdAB* (group B); and mutations that co-occur at similar frequencies with GyrA-S83X and *ccdAB* (group C). Within group C, we see mutations with an intermediate level of co-occurrence with GyrA-S83X and CcdB (group C1) mutations with a high level of co-occurrence with these two genes (group C2).

Group A includes all QRDR mutations, except for ParC-E84X. This is expected given the “gateway” status of GyrA S83X for QRDR trajectories. The E84X exception is interesting and is discussed in detail below. Group B includes most non-QRDR mutations, which is consistent with the idea that CcdB is acting as a positive selection for this group of mutations. The stark difference between the two groups implies that one or multiple QRDR mutations (given the high co-ocurrence of QRDR mutations, the contribution of individual mutations is hard to establish) are likely negatively epistatic with the presence of the *ccdAB* operon.

Group C1 includes two single QRDR mutations (GyrB-S492N, GyrB-A618T) that likely represent mutations without functional interactions with either GyrA-S83X or *ccdAB*.

Group C2 includes ParC-E84X, the combination [S57T D197E K200N D309E L344R] and the triplet [A192V A471G Q481X]. This group of mutations could represent permissive mutations that allow GyrA-S83X and *ccdAB* to co-exist stably. Indeed, if QRDR mutations exhibit negative epistasis with *ccdAB*, quinolone treatment would be expected to drive the selection of permissive mutations that override the negative epistatic interaction. Consistent with this idea, we do see two sets of these putative permissive mutations (E84V, and [A192V,A471G,D475E,Q481H]) in almost all ST131 samples. It is tempting to speculate that the co-existence of QRDR and non-QRDR mutations may provide some fitness advantage that could have contributed to the remarkable epidemic expansion of the ST13-H30 clone.

Interestingly, group C2 mutations are all found in ParC, the same gene that carries the ParC-D475E gateway mutation for non-QRDR trajectories. Consistent with the idea that mutations in group C2 represent compensatory mutations, the specific amino acid substitution at one QRDR position determines the amino acid substitution at the proposed compensatory site. In the presence of the unusual GyrA-S83W and ParC-S80R mutations, we found an unusual substitution in ParC (E84Q) These observations imply that topoisomerase IV may be a secondary target for the toxin CcdB in *E. coli*. Alternatively, these “permissive” mutations in the “A” subunit of topoisomerase IV may be enhancing fitness indirectly through changes in global supercoiling, as previously suggested for *gyrA* mutations in *Campylobacter jejuni* (36) and for *gyrA* and *parC* mutations in *Salmonella typhi* (37).

### Truncated CcdB sequences support mutual exclusion between non-QRDR topoisomerase II mutations and GyrA-S83X

While the positive association between most non-QRDR mutations and *ccdAB* and the negative association between *ccdAB* and QRDR mutations are intriguing, firm conclusions cannot be drawn because the presence of *ccdAB* and non-QRDR mutations is restricted to a sector of the cladogram **(Fig. 3**). Nonetheless, we observed that the *ccdB* sequence is often truncated. Of the 52 STs having the CcdB sequence, 19 have a sequence that is predicted to translate into a protein of between 95 and 40 amino acids long instead of the full 105 amino acids. In the cladogram, we see only 9 transitions between full-size and truncated forms of CcdB, but these transitions are seen in all major phylotypes (F, D and B2), indicating that, while very stable, a switch from full-size to truncated CcdB has occurred multiple times independently (**Fig. 3**, panels 15-16).

Frequent truncations can reflect genetic drift or a negative selection. We hypothesized that if the observed truncations reflect a negative selection, and if this negative selection is driven by CcdB’s incompatibility with QRDR mutations, the distribution of QRDR mutations in CcdB-truncated STs should be more similar to the distribution of these mutations seen in CcdB-less strains. This would address the bias in the distribution of non-QRDR mutations and *ccdAB* in the cladogram. Also, given that truncations are likely to affect the function of CcdB, this analysis takes us closer to a causal link between the presence of CcdB and a negative impact on the fitness of QRDR mutant cells.

The results of this analysis, shown in **Fig. 6a** and **Fig. S5**, support the idea that CcdB truncations are the result of a negative selection in the presence of QRDR mutations. We see that in CcdB-truncated STs, the fraction containing GyrA-S83X mutations is much higher than in STs with full-length CcdB. We also hypothesized that in samples with CcdB-truncated STs, we would see a decrease in the representation of some non-QRDR mutations, similar to what we see in CcdB-STs. Indeed, we do see this effect in two mutations: GyrA-A828X and ParE-V136I, and more moderately also in GyrB-E185D and GyrB-A618T (**Fig. 6b**). These observations support the idea that CcdB is positively selecting for most non-QRDR mutations.

**Fig. 6.**
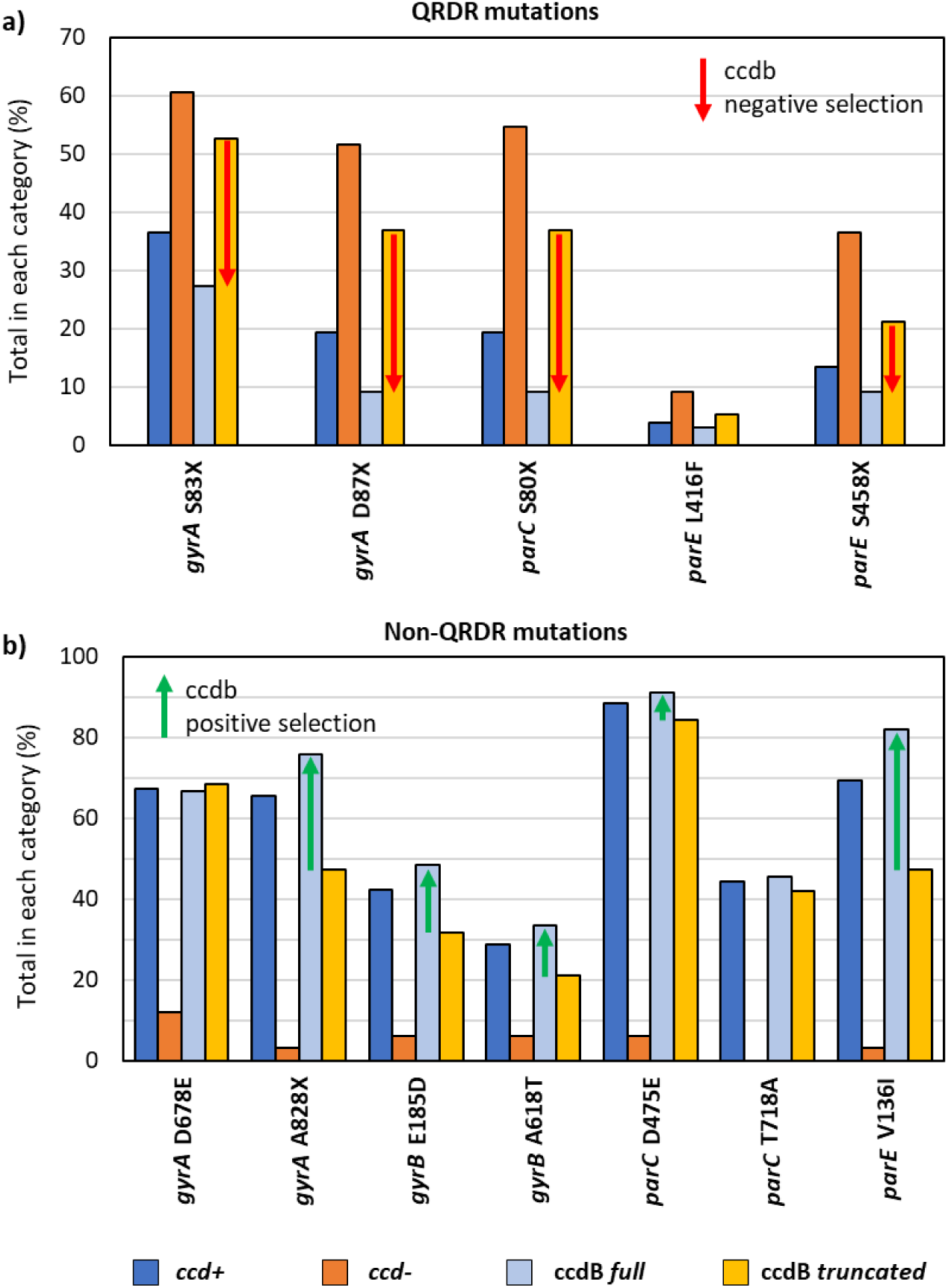
Impact of ccdAB status on the representation of selected QRDR and non-QRDR mutations. Fraction of STs containing ccdAB+, ccdbAB-, ccdbAB+ full-length and ccdAB+ truncated sequences for the individual mutant positions listed in the x-axis. Differences that are consistent between ccdAB- and ccdAB truncated are highlighted with arrrows **a**. The fraction shown corresponds to five representative QRDR mutations, listed in the x-axis. **b**. The fraction shown corresponds to seven representative non-QRDR mutations

Overall, these observations confirm the negative genetic interaction between CcdB and/or non-QRDR mutations and QRDR mutations already suggested by our Jaccard Index and phylogenetic analyses. They also suggest that CcdB is likely represents a positive selection driving the fixation of most of the non-QRDR mutations that we have identified, although the degree of selection varies between individual mutations.

### Impact of QRDR and non-QRDR mutations on antibiotic resistance

To determine the contribution of CcdB-selected non-QRDR mutations on antibiotic resistance, we trained a variety of logistic regression models to predict the fluoroquinolone sensitivity or resistance of individual strains using three of these mutations and chromosomal ccdB as features. As positive controls, we also included known determinants of resistance as features in our model of fluoroquinolone resistance; as a negative control, we included the plasmid form of ccdB.

Given that gyrase and topoisomerase IV are both encoded in the chromosome, their genetic variance is expected to be influenced by the underlying population structure (38). To estimate a baseline representing the generic contribution of population structure to the model, we incorporated three population structure descriptors (PC1, PC2, PC3) as additional continuous variables. These descriptors were obtained through a multi-dimensional scaling (MDS) projection of individual single nucleotide polymorphism (SNPs) variance across all genomes in the dataset, an approach previously reported as a control for regression models involving bacterial genomic data (see **Methods** and **Supplementary Material**) (39,40).

To prevent overfitting, all our logistic regression models were regularized using an *I2* penalty (see methods) and we performed 5-fold cross-validation of sequence types across 1,000 iterations. The predictive power of classification models can be measured using the area under the ROC curve (AUC), which is a value between 0.5 (no better than random) and 1.0 (perfect accuracy and recall). As a control, we randomly permuted the response variable and obtained the expected mean AUC value of 0.5 (see **Methods**).

The results of this modeling are summarized in **Table 2**. They show that population structure descriptors can only modestly predict fluoroquinolone resistance (AUC=0.589). The incorporation of additional regressors dramatically increases the predictive value of the models (AUC > 0.87 for models 2 to 7 in **Table 2**). Note that the inclusion upregulation of drug efflux, which is found in about 50% of fluoroquinolone resistant UTI strains (41), as a feature is not necessary in our dataset for the construction of a highly accurate predictive model.

**Table 2.** Summary of different logistic regression classifier models trained to predict the fluoroquinolone resistance of individual *E. coli* strains using genotypic features. The table collects the average coefficients associated to different features (β_i_) and the figures of merit in the test set for the different models. See methods for details.

|  | Model 1 | Model 2 | Model 3 | Model 4 | Model 5 | Model 6 | Model 7 |
| --- | --- | --- | --- | --- | --- | --- | --- |
| **GyrA_S83X** | - | 0.826 | 0.821 | 0.809 | 0.807 | 0.772 | 0.765 |
| **GyrA_D87X** | - | 1.067 | 1.066 | 1.057 | 1.054 | 0.992 | 0.991 |
| **ParC_D475E** | - | - | -0.143 | -0.130 | -0.120 | - | -0.160 |
| **ParE_V136I** | - | - | - | -0.177 | -0.174 | - | - |
| **GyrA_A828X** | - | - | - | - | 0.019 | - | - |
| **CcdB chromosome** | - | -0.117 | -0.086 | - | -0.070 | - | - |
| **CcdB plasmid** | - | 0.215 | 0.221 | 0.224 | 0.225 | 0.170 | 0.177 |
| **QnrS** | - | 0.291 | 0.292 | 0.286 | 0.285 | 0.265 | 0.265 |
| **Aac(6')-Ib-cr** | - | 0.380 | 0.378 | 0.380 | 0.377 | 0.347 | 0.343 |
| **PC1** | 0.215 | 0.039 | 0.058 | 0.079 | 0.082 | -0.044 | -0.020 |
| **PC2** | 0.635 | 0.252 | 0.247 | 0.240 | 0.238 | 0.215 | 0.208 |
| **PC3** | -0.001 | 0.011 | 0.008 | 0.013 | 0.011 | 0.003 | -0.000 |
| **CcdB ch. cluster 1** | - | - | - | - | - | -0.267 | -0.249 |
| **CcdB ch. cluster 2** | - | - | - | - | - | 0.316 | 0.329 |
| **CcdB ch. cluster 3** | - | - | - | - | - | -0.180 | -0.169 |
| **AUC** | 0.593 | 0.876 | 0.877 | 0.876 | 0.874 | 0.884 | 0.889 |
| **Precision** | 0.300 | 0.891 | 0.883 | 0.880 | 0.878 | 0.899 | 0.890 |
| **Recall** | 0.188 | 0.834 | 0.825 | 0.757 | 0.738 | 0.842 | 0.835 |

According to the average β_i_ coefficients (listed on **Table 2**), the classification performance is largely driven by the two most abundant QRDR mutations included in our model (GyrA S83X and D87X). Note that the β_i_ coefficient for GyrA D87X is superior to that of GyrA S83X in our four models (ca. 1.0 and 0.8, respectively). Given that all GyrA D87N mutants also have the GyrA S83X mutation (**Fig. 1a**), the implication is that the GyrA-S83X mutation by itself may not be sufficient to confer fluoroquinolone resistance in standard clinical assays. A previous study reached a similar conclusion (42), helping validate our approach.

The plasmid markers *qnrS* and *aac(6)-lb-cr* make more modest contributions to the resistance classification by the models (β_i_ coefficients in the 0.28-0.38 range). *aac(6)-lb-cr* always co-occurs with the gateway GyrA-S83X mutation, consistent with the idea that it represents a downstream acquisition enhancing QRDR mutation-mediated resistance rather than a mechanism that can confer high levels of fluoroquinolone resistance on its own (43). Also note that the vast majority of the plasmids found in our dataset were IncF plasmids (**Dataset S1**), and IncF plasmids are known to be the main carriers of *aac(6)-lb-cr* (43,44).

Plasmid *ccdAB* also makes a modestly positive contribution to the model (β_i_ coefficient in the 0.21-0.23 range) and this is again likely due to the high prevalence of *ccdAB* in incF plasmids, where it enhances plasmid stability by coupling host cell division to plasmid proliferation (45).

We included three non-QRDR mutations in the models. The two that are most upstream in the trajectory described in **Fig. 1b (**ParC-D475E and ParE-V136I) are negatively correlated with fluoroquinolone resistance (β_i_ coefficient between −0.12 and −0.17). This is consistent with very low JI coefficients for pairs involving QRDR and non-QRDR mutations in **Fig. 2**, and with the trend toward mutual exclusion shown in **Fig. 5**. Similarly, the presence of chromosomal *ccdAB* has a modest negative contribution to the model, consistent with the partial mutual exclusion of QRDR mutations shown in **Fig. 5** and **Fig. 6**. The more downstream mutation (GyrA-A828X) is neutral.

Importantly, when we discriminate between the full-length (cluster 1), moderate truncation (cluster 2) and severe truncation of CcdB (cluster 3), the sign turns more negative for the full-length (−0.26) and flips sign for the moderate truncation (+0.32), again pointing to the causality between the presence of full-length CcdB and decreased fitness in the presence of QRDR mutations and/or fluoroquinolone.

## Conclusions

We provide strong genetic evidence suggesting that the presence of the chromosomal form of the toxin CcdB is driving the selection of a set of non-QRDR mutations in type IIA topoisomerases. The toxin CcdB is known to target DNA gyrase, but most previous studies focused on plasmid-borne *ccdAB* genes (35,46). The biological role of chromosomal TA systems is largely unknown (33,34), although a role in biofilm formation and antibiotic persistence has been suggested (47,48). The co-evolution of type II topoisomerase genes with chromosomal *ccdAB* implies that, like its plasmid counterpart, chromosomal CcdB targets gyrase. This is in agreement with recent reports (49).

In addition to mutations in gyrase, the non-QRDR mutations identified to be under CcdB selection involve the two subunits of topoisomerase IV. We also find “permissive” mutations that appear to facilitate the co-existence of QRDR and non-QRDR mutations, which are all found in the ParC subunit of topoisomerase IV. All these observations suggest that topoisomerase IV is likely a target for the chromosomal form of CcdB in *E. coli*. In this scenario, CcdB would parallel quinolone drugs (5,8-10) and qnr resistance factors (17-20), which target both gyrase and topoisomerase IV. Alternatively, mutations in topoisomerase IV may represent compensatory mutations altering global supercoiling in response to CcdB-mediated inhibition of gyrase.

The early acquisition of the chromosomal *ccdAB* operon in the evolution of phylotypes D, F, U, and B2 and its stable maintenance suggests that the interaction of CcdB and type II topoisomerases has meaningful biological consequences. Note that (unlike its plasmid counterpart) chromosomal CcbB is largely bacteriostatic and therefore can be better tolerated by the cell (49). Our analysis also suggests a mutual exclusion between CcdB and QRDR mutation trajectories (**Fig. 2**,**5**,**6**). The nature of this is antagonism is unclear. The binding of free CcdB toxin to mutant topoisomerases may decrease their activity to the point that it is insufficient for the optimal growth of the cell. Alternatively, the antagonism could be due to negative epistatic interactions between QRDR and non-QRDR mutations under ccdAB selective pressure. The present work lays the foundation for the experimental verification of these hypotheses.

This work also has translational implications. The observed antagonism between cc*dAB* and QRDR mutations suggests that the presence of full-sized *CcdB* and permissive mutations in ParC can be used as genetic markers for the risk of evolution of quinolone resistance through the acquisition of QRDR mutations. This is relevant to epidemiological surveillance and implies that certain strains may be genetically more refractory to evolving resistance through mutations in topoisomerase. Also, the potential for free CcdB and non-QRDR mutations to modulate sensitivity to quinolones needs to be explored.

## Methods

### Sample and data collection

The first dataset (UW233) contains 233 draft genomes from a published study conducted at the University of Washington (24). This study represents an unbiased sampling of clinical extraintestinal *E. coli* isolates from patients suffering from sepsis or from urinary tract at the University of Washington Medical Center. The methods for antibiotic susceptibility profiling are described in (24). These samples had a relatively low rate of ciprofloxacin resistance (27%).

To increase the density of ciprofloxacin-resistant isolates, we generated a second dataset, DHMMC119, consisting of 119 draft genomes from a collection of extended-spectrum beta-lactamase (ESBL) resistant isolates obtained from the Dignity Health Mercy Medical Center in Merced, CA, USA. We reasoned that fluoroquinolone resistance and ESBL resistance are frequently linked (50) and indeed, the fraction of ciprofloxacin resistant strains in this dataset was 94%.

Clinical samples were collected from patients with respiratory, blood (wound), or urinary tract infections at Dignity Health Mercy Medical Center (DHMMC) in Merced, California, between June 2013 and August 2015. Isolates were identified and tested for ESBL-mediated resistance using Vitek 2 Version 06.01, and were also tested for susceptibility against 16 antibiotics using broth micro-dilution minimum inhibitory concentration (MIC) testing. The isolates were initially categorized as Resistant (R), Intermediate (I), or Susceptible (S), based on the MIC Interpretation Guideline – CLSI M100-S26 (2015). For the logistic regression models, isolates displaying Intermediate resistance were considered to be Resistant to generate an approximate binary classification.

### Sequencing, assembly, and quality control of ExPEC genomes

For the samples from a previous study conducted at the University of Washington, all 384 ExPEC draft genome assemblies were downloaded from the NCBI database, with accessions in the range JSFQ00000000–JSST00000000.

For the DHMMC isolates, the whole-genome sequencing was performed using the MiSeq and HiSeq paired-end (2×250 bp) sequencing technologies with TruSeq DNA library preparation. Twenty-four MiSeq and 110 HiSeq libraries of paired-end reads were obtained and assembled using SPAdes v3.5.0 (51) with read error correction by BWA-spades, and with automatic *k*-mer size selection. Additional details of the DNA extraction, sequencing, and assembly procedures are provided in **Supplementary Material**.

Each DHMMC and UW assemblies were assessed for completeness based on the presence of 143 protein-coding genes considered to be essential for normal growth of *E. coli* MG1655 (Set A in Supplementary Table 2 from (52)). To achieve consistent assembly qualities across the DHMMC and UW datasets, assemblies containing fewer than 126 full-length essential genes were removed. Remaining were 119 DHMMC and 223 UW assemblies, which are jointly referred to as USWest352.

### Phylogenetic classification of strains

Phylogenetic classification of the 352 draft genome assemblies of USWest352 was performed using three methods that provided increasing levels of resolution: the EzClermont phylotyping method (53), the Achtman multi-locus sequence typing (ST) method (25), and based on nucleotide-level variation across whole-genomes. In the latter, an all-by-all pairwise distance matrix was constructed, in which the genetic distance between each pairing of genomes was estimated as the number of nucleotide positions varying between the two genomes. Variant calling for each genome was performed using the *nesoni consensus* script from the nesoni package v0.128. The distance matrix was generated using the *nesoni nway* script from the nesoni package, specifying *E. coli* EC958 as the reference genome (27). A neighbor-joining tree was constructed based on the distance matrix using SplitsTree4 (54).

### Detection of CcdB and ParE genes

Sequences for the chromosomal CcdA/CcdB and ParE toxin and antitoxin genes were retrieved from TADB2 (58). These sequences were then queried against the USWest352 database of bacterial assemblies, indexed using NCBI blast-2.10.1+ (56). Megablast settings were used, requesting a sequence identity above 85% and an alignment length ≥ 200 bases. Strains were considered positive for the ParE or CcdA/CcdB toxin-antitoxin if either the toxin or the antitoxin were detected. Notably, in all the 281 strains in which CcdB toxin was detected, the corresponding CcdA antitoxin was also detected.

### Detection of Qnr genes

A variety of Qnr genes have been reported in the literature (55). We compiled a custom list of Qnr DNA sequences including QnrA1, QnrA2, QnrA3, QnrA4, QnrA5, QnrA6, QnrB1, QnrB2, QnrB3, QnrB4, QnrB5, QnrB6, QnrB7, QnrB8, QnrB9, QnrB10, QnrB11, QnrB12, QnrB13, QnrB14, QnrB15, QnrB16, QnrB17, QnrB18, QnrB19, QnrS1, QnrS2, and QnrS3. We then queried this custom list against the indexed USWest352 database using NCBI blast-2.10.1+ (56). Megablast settings were used, requesting a sequence identity above 90% and an alignment length ≥ 200 bases. Each strain was the labeled as “No Qnr”, “QnrA”, “QnrB”, or “QnrS” depending on the Qnr family detected.

### Jaccard index calculations

The Jaccard index (JI) measures the co-occurrence of two mutations (or other binary features), and ranges from 0 (mutual exclusion) to 1 (total co-occurrence). To calculate this index, we used the following formula:

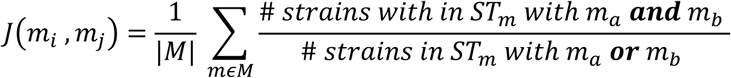

where *M* is the set of Achtman STs in which mutations m_a_ or m_b_ have been observed.

### Intra-ST spread index calculations

We define a ST-weighted spread index for mutation m_i_ as follows:

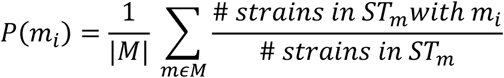

where *M* is the set of Achtman STs in which mutation m_i_ had been observed in three or more strains.

### Relative abundance of S83X and CcdB in different STs

To normalize the relative abundance on a ST basis, we use the following weighted average expressions (fraction indices, F). The ST-weighed fraction of mutants m_i_ being S83X-positive would be:

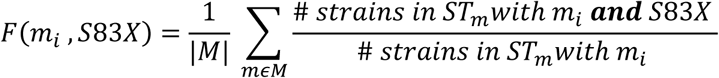

where *M* is the set of Achtman STs in which mutation m_i_ had been observed.

Similarly, the ST-weighed fraction of mutants m_i_ being CcdB-positive would be:

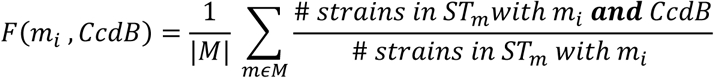

where *M* is the set of Achtman STs in which mutation m_i_ had been observed.

### Location of CcdA/CcdB genes

The location (chromosome vs. plasmid location) of the CcdA/CcdB TA systems was investigated using two methods. First, the co-occurrence of either CcdA or CcdB and a replicon sequence in the same contig was evaluated. A database of replicon sequences was compiled by copying the curated database of the software PlasmidFinder (59). The PlasmidFinder database currently includes 133 unique plasmid replicon sequences. The replicon sequences were queried against the USWest352 database using megablast with a sequence identity above 85% and an alignment length ≥ 100 bases. 3,950 replicons could be detected throughout the USWest352 database, the majority of them from the *Enterobacteriaceae* sub-database. The total number of replicons detected for each ST is shown in **Fig. S4**.

The most common replicons belong to the group IncFII. However, none of the replicons detected were found in contigs where the CcdA or CcdB sequences had been detected.

On the other hand, we applied the software mlplasmids (60) to the specific contigs were CcdA or CcdB had been identified. This software considers the distribution of pentamer sequences in a contig as input to a machine-learning model, which classifies the contig as chromosomic or plasmidic. We used the settings for *Escherichia coli*. For all CcdA/CcdB-containing contigs, the software assigns a >93% probability that the contig is chromosomic (for reference, the default threshold in the software is 50% probability).

### CcdB polymorphisms

The CcdB sequences were looked at in more detail. The aligned subject sequences from Megablast were translated with EMBOSS Transeq tool using the bacterial codon table https://www.ebi.ac.uk/Tools/st/emboss_transeq/. To help select the correct reading frame, ExPASy translate was referred to for guidance https://web.expasy.org/translate/. Once the DNA sequences were translated to amino acid sequences, they were all aligned to the amino acid sequence predicted for the reference CcdB using blastp default settings (56). The goal of this blastp is to align the protein sequences. The results were then used to estimate the length of the CcdB protein, determined by the alignment length or the position of the first stop codon. No CcdB sequence was cut to being located in one end of the contig. The CcdB sequences were then clustered in three groups depending on their length: cluster 1 (full-length, 105 amino acids), cluster 2 (partially truncated, 95 amino acids), cluster 3 (40-84 aligned amino acids).

### Determination of marker predictive values using logistic regression

A variety of logistic regression models were trained to predict the fluoroquinolone sensitivity or resistance of individual strains using different selected genetic features. These features indicate the presence (1) or absence (0) of individual genes or mutations. The response variable was the binary fluoroquinolone resistance, as indicated in the Supporting Dataset S1.

The logistic regression classifier takes the form:

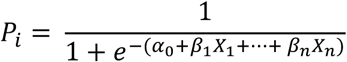

where *p*_*i*_ is the probability that strain *i* is fluoroquinolone resistant, *p*_*i*_ ∈ [0,1]; α_0_ is the intercept, β_1_ … β_n_ are the coefficients associated to the features X_1_ … X_n_. A positive coefficient increases the probability that a strain be classified as resistant if it has the corresponding feature, whereas a negative coefficient lowers the probability that a strain be classified as resistant if it has the feature.

To account for the way genetic variance is structured in the population, we incorporated to the models three population structure descriptors (PC1, PC2, PC3) as additional continuous variables. These were obtained through a multi-dimensional scaling (MDS) projection of individual single nucleotide polymorphisms (SNPs) across all genomes in the dataset. We performed classical multi-dimensional scaling (cmdscale package in R) on the all-by-all (*n*×*n, n* = 352) nucleotide distance matrix generated using the nesoni package as described above. Three components were thus included as regressors in the model.

Furthermore, we regularized the logistic regressor models using an *l2* penalty, with a regularization coefficient *C*=0.05 in all cases, as implemented in scikit-learn 0.24.1.

80% of the sequence types in the dataset were randomly assigned to the training set, and the remaining 20% were assigned to the test set. This splitting in terms of sequence types instead of individual strains is meant to provide a more stringent test of the predictive power of the models on more distant strains. To obtain more robust estimates, this process was repeated 1,000 times for each model, and the average of the coefficients and figures of merit was computed.

## Abbreviations

AUC: Area Under the Curve
JI: Jaccard Index
MDS: Multi-Dimensional Scaling
QRDR: Quinolone Resistance-Determining Region
ROC: Receiver Operating Characteristic
SNP: Single Nucleotide Polymorphism
ST: Multi-locus Sequence Type
TAS: Toxin-Antitoxin System
UTIs: Urinary Tract Infections

## Acknowledgements

The authors acknowledge Dr. Steve Salipante (University of Washington) for his help processing and interpreting the UW sequences and for generously sharing 39 strains of his collection, Lily Tran for her initial assistance processing the data, Caison Warner for his assistance with the genomic mapping of *ccdAB*, and the DHMMC for their partnership in the collection of the DHMMC119 set of clinical samples. This work was funded by CITRIS Seed Funding proposal 2015-324 and by NIAID award 1R41AI122740-01A1 to MC and MB in partnership with Maverix Genomics.

## Notes

### Competing Interest Statement

The authors have declared no competing interest.

