## Supplemental Figures and Tables for "Genetic interplay between type II topoisomerase enzymes and chromosomal *ccdAB* toxin-antitoxin in *E. coli*"

**Table S1** Summary of the strain composition of the datasets.

| Dataset | Size | Source | Sampling enrichment | Total STs | Top 5 STs |
|---|---|---|---|---|---|
| UW233 | 233 | University of Washington | none | 75 | ST131 (15.5%), ST95 (11.6%), ST73 (9.4%), ST127 (6.4%), ST69 (5.2%) |
| DHMMC119 | 119 | Dignity Health Mercy Medical Center | ESBL-producers | 22 | ST131 (58.0%), ST648 (10.9%), ST44 (5.0%), ST38 (4.2%), ST405 (3.4%) |
| USWest352 | 84 | UW233, DHMMC119 | ESBL-producers (partial) | 84 | ST131 (29.8%), ST95 (8.0%), ST73 (6.3%), ST127 (4.8%), ST69 (3.7%) |

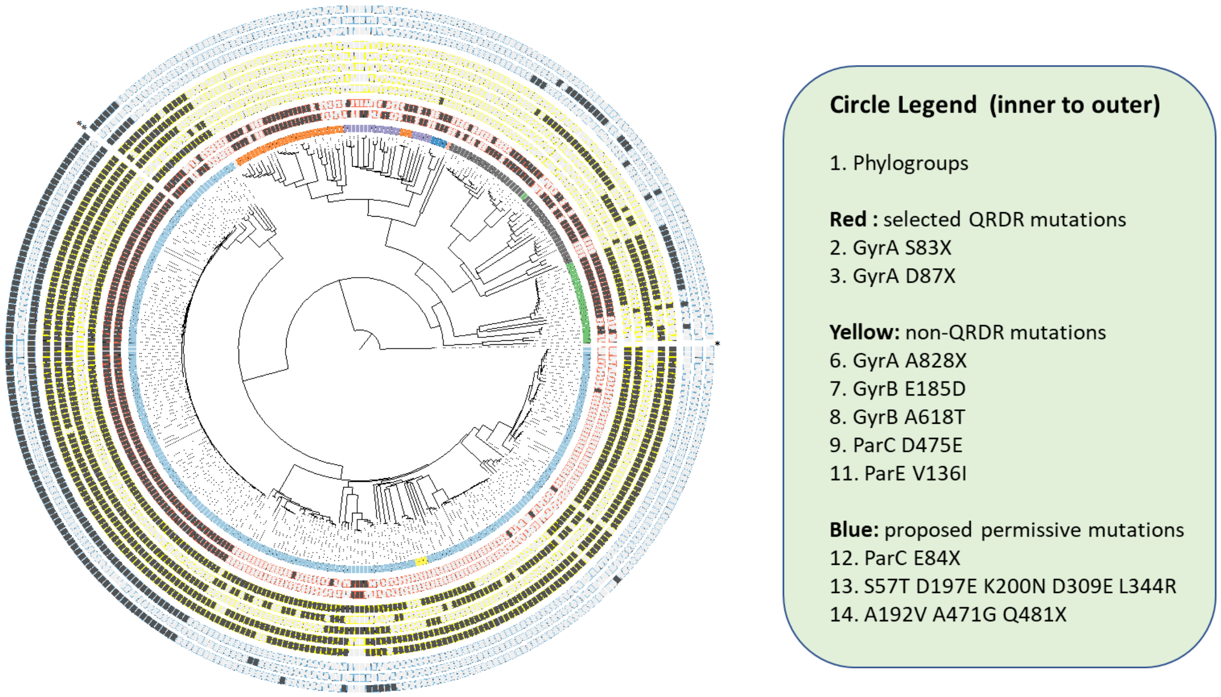


**Fig. S1** Visualization of the phylogenetic distance between 352 extraintestinal E. coli isolates obtained from two hospitals in the U.S. West Coast. The phylogenetic relationship between the samples is represented as a neighbor-joining tree, computed based on pairwise comparisons of 537,420 genomic SNPs. The squares represent the presence of selected mutations or genes. Panel 1: Phylotype A (orange), B1 (purple), B2 (light blue), C (dark blue), D (dark grey), E (light red), F (green), U (yellow). Panels 2-3 (outlined in blue) Selected QRDR mutations GyrA-S83X and -D87X. Panels 4-8 (outlined in yellow) Selected non-QRDR mutations: GyrA A828X, GyrB E185D and A618T, ParC 475E, and ParE V136I . Panels 9-11 (outlined in blue): Proposed permissive mutations ParC: ParC E84X, [S57T D197E K200N D309E L344R], and [A192V A471G Q481X] .

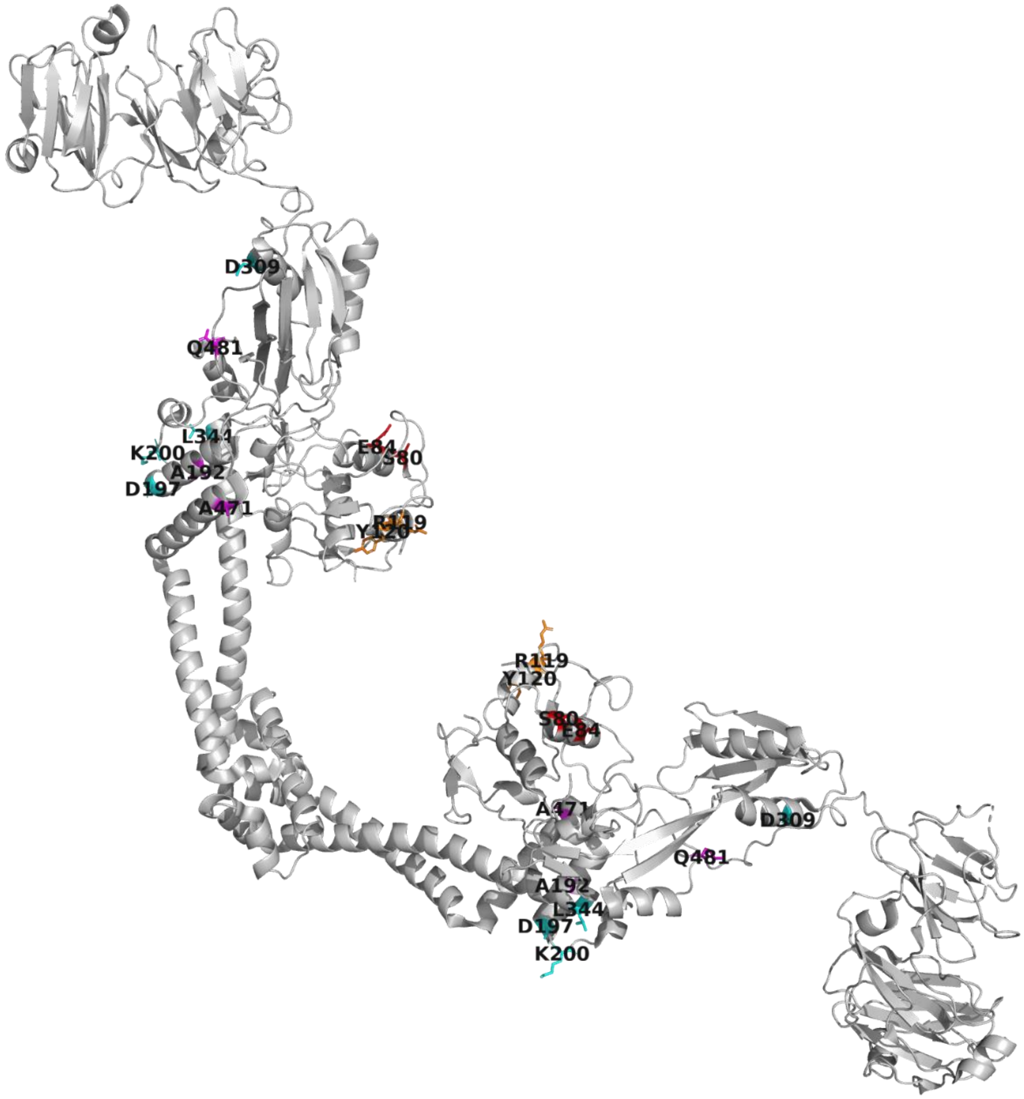


**Fig. S2** Location of proposed permissive mutations in the dimeric structure of ParC: [D197 K200 D309 L344] in cyan, and [A192V A471G Q481X] in magenta. For reference, QRDR residues S80 and E84 are shown in red, whereas R119 and Y120 in the active site are shown in orange. Interestingly, the putative permissive mutations except E84X (red) fall on the opposite side of the active site. Figure prepared using PyMOL from PDB model 1zvu.

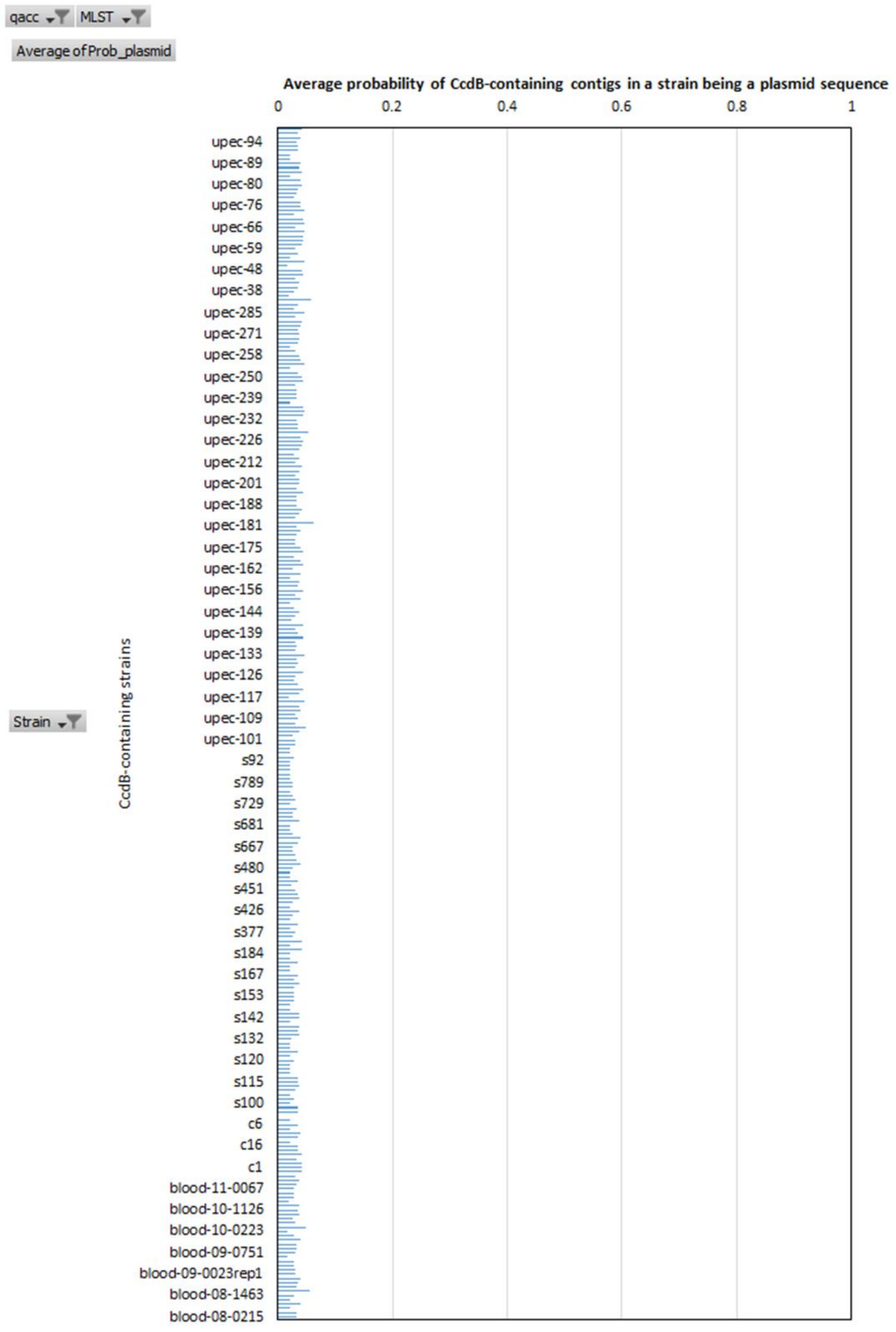


**Fig. S3** Plasmid classification probability by the machine-learning model *mlplasmids* for those contigs containing the chromosomal CcdB for each strain in USWest352. In all cases, the classifier is highly confident that the CcdB gene studied is located in the chromosome, assigning a very low probability that the contigs containing these genes are plasmidic.

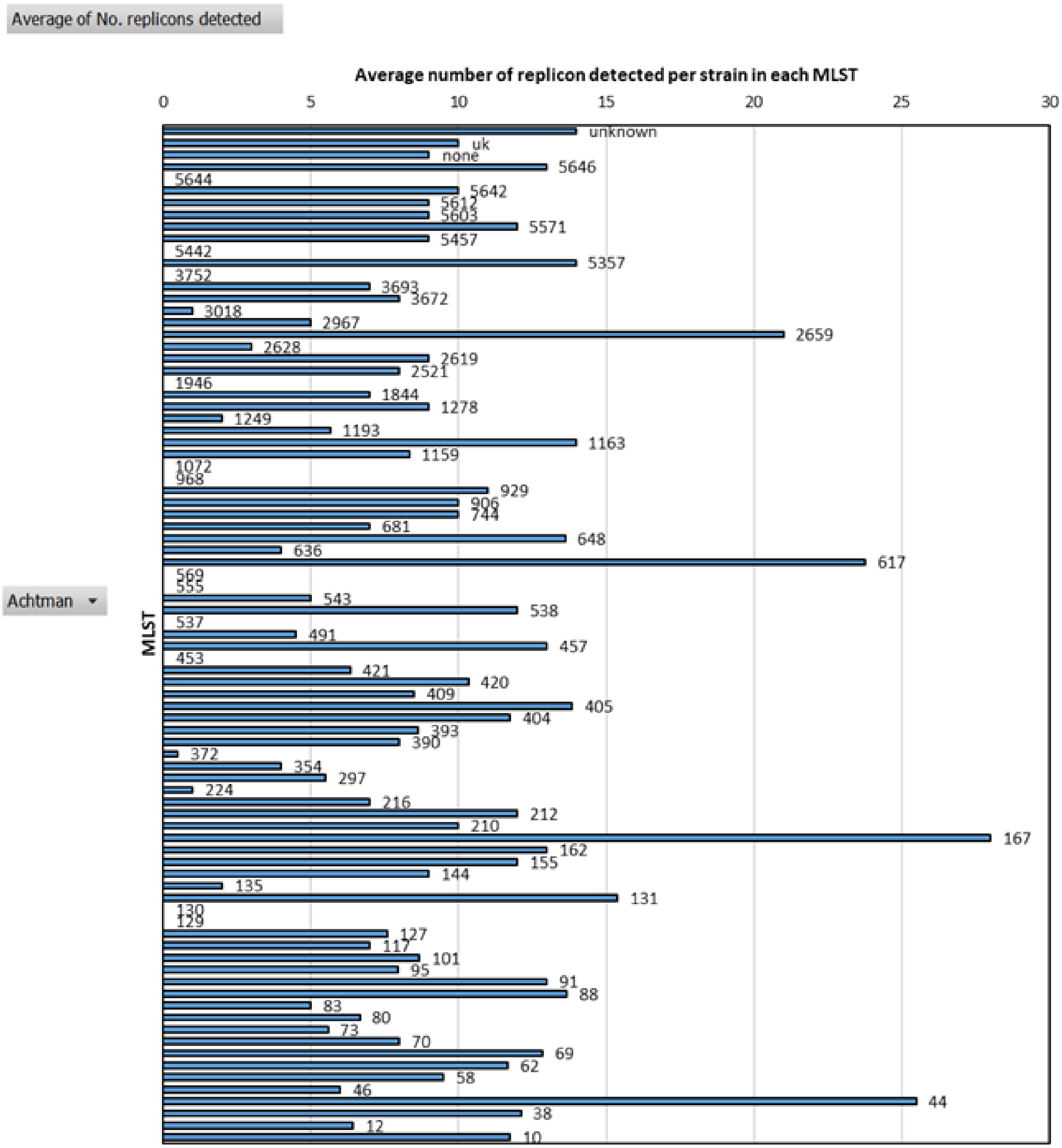


**Fig. S4** Average number of replicons detected in the assemblies of the different STs in USWest352. A wide variation in the number of replicons detected is observed across different STs, from none to over 20 in some STs. Most STs displayed between 5 and 10 replicons, which is consistent with the distribution of replicons in completely assembled E. coli genomic sequences deposited in NCBI (61).

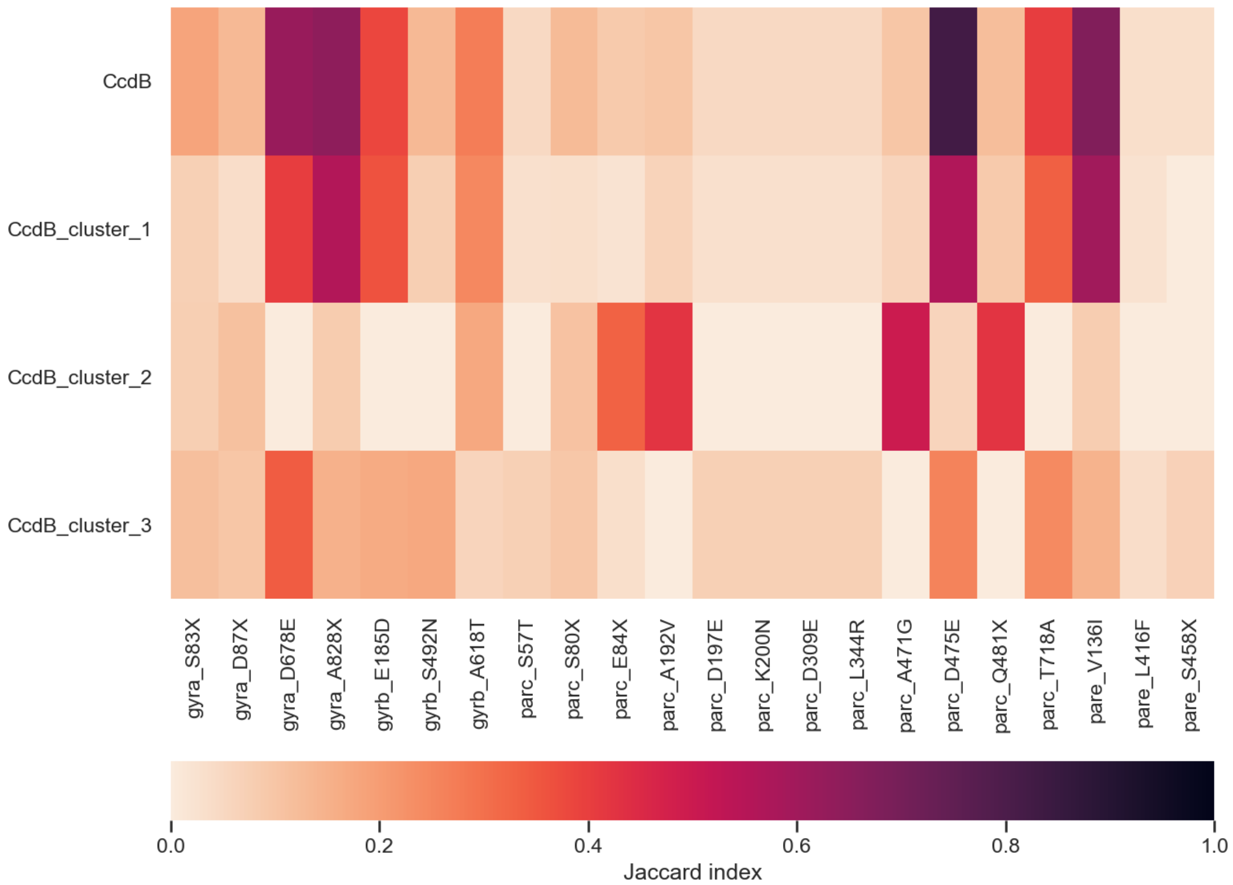


**Fig. S5** Jaccard indices between different CcdB variants and some common mutations in type II topoisomerases. The CcdB variants are clustered according to their predicted length upon expression: cluster 1 (full-length, 105 amino acids), cluster 2 (partially truncated, 95 amino acids), cluster 3 (40-84 aligned amino acids). A strong co-occurrence is observed for the full-length variant and some mutations in GyrA (D678E, A828X) and ParC (D475E and V136I). This co-occurrence is greatly weakened for the truncated variants. The Jaccard indices are weighted by MLST, as indicated in the methods.
